# Alpha Protein Kinase 3 Gene Therapy Restores Heart Function in Mouse and Human Models of Cardiomyopathy

**DOI:** 10.1101/2025.07.31.667858

**Authors:** James W. McNamara, Ellen B. Keen, Rebecca Sutton, Yinghan She, Neda R. Mehdiabadi, Brendan Griffen, Richard M. Mills, James E. Hudson, Drew M. Titmarsh, Enzo R. Porrello, David A. Elliott

**Author notes:** Corresponding Authors: James W. McNamara, Enzo R. Porrello and David A. Elliott., Murdoch Children’s Research Institute, 50 Flemington Road, Parkville, Melbourne, VIC, 3052, Australia. Contributed Equally (Enzo R. Porrello, David A. Elliott).

## Abstract

Truncating variants in the Alpha Kinase 3 (ALPK3) gene have recently emerged as an important cause of genetic cardiomyopathy globally and here we demonstrate the efficacy and safety of a viral-based gene replacement therapy for ALPK3. Given the loss-of-function nature of these variants, we reasoned that a gene replacement approach would improve heart function in this patient population. We demonstrate that the delivery of full length ALPK3 via adeno-associated virus could restore contractile function in human cardiac organoids and in vivo mouse models carrying clinically relevant mutations in ALPK3. The role of disrupted proteostasis networks in multiple forms of genetic cardiomyopathy suggest that delivery this novel AAV-ALPK3 may provide functional benefit outside of cardiomyopathy induced by ALPK3. Titin truncating variants (TTNtv) are the most common cause of dilated cardiomyopathy, and interestingly also contributes to the M-Band protein quality control network coordinated by ALPK3. Notably, in human cardiac organoids carrying a TTNtv we observed that the ALPK3 gene therapy could completely restore contractile deficits. This opens the exciting prospect for indication expansion of AAV-ALPK3 into other forms of cardiomyopathy that currently have no therapeutic options.

## Main

Advances in our understanding of cardiomyopathy genetics have led to an emerging wave of precision therapies targeting the cause of disease, rather than management of secondary symptoms.^1^ The success of the myosin inhibitor mavacamten provides a striking example of how targeting the underlying mechanisms driving disease can improve clinical outcomes for patients with cardiomyopathy.^1^ Several gene replacement therapies are showing promise in clinical trials for some of the most common inherited forms of hypertrophic and arrhythmogenic cardiomyopathies caused by mutations including *MYBPC3* and *PKP2*.^1^ Therefore, expansion of gene therapies to other inherited cardiomyopathies may represent an opportunity to provide life-changing treatment to patients.

Loss of function variants in the myogenic Alpha Protein Kinase 3 (*ALPK3*) gene have emerged as an important genetic cause of cardiomyopathy, accounting for around 2% of hypertrophic cardiomyopathy cases^2,3^. Biallelic variants cause a severe, and often lethal, form of neonatal cardiomyopathy,^2,3^ while autosomal dominant variants are strongly linked to adult onset hypertrophic (and to a lesser extent dilated and arrhythmogenic) cardiomyopathy.^4,5^ Importantly, ALPK3 orchestrates a protein quality control hub at the sarcomere to maintain turnover of contractile proteins.^6-8^ Given the loss of function nature of *ALPK3* variants, we hypothesised that gene replacement therapy may provide functional benefit for *ALPK3* cardiomyopathy.

To evaluate this therapeutic approach, we differentiated a previously characterised^6^ human pluripotent stem cell (hPSC) line carrying a loss of function mutation in *ALPK3 (ALPK3*^mut^) into cardiac monolayers using established methodology (approved by Royal Children’s Hospital Research Ethics Committee (HREC) 33001A and 93025).^9^ The *ALPK3*^mut^ line exhibits severe sarcomere disorganisation due to ALPK3 deficiency (Figure 1A and ^6^). *ALPK3*^mut^ cardiac cells were transduced with an adeno-associated viral (AAV) vector, serotype 6, encoding a full length *ALPK3* gene with a C-terminal 3xFLAG epitope. The FLAG epitope localized correctly to the sarcomeric M-band, confirming proper translation and trafficking of full length ALPK3 (Figure 1A). Importantly, AAV-mediated *ALPK3* expression dramatically restored the severe sarcomeric disorganisation caused by loss of *ALPK3*^6^, as evidenced by the restoration of ACTN2 organisation (Figure 1A).

**Figure 1.**
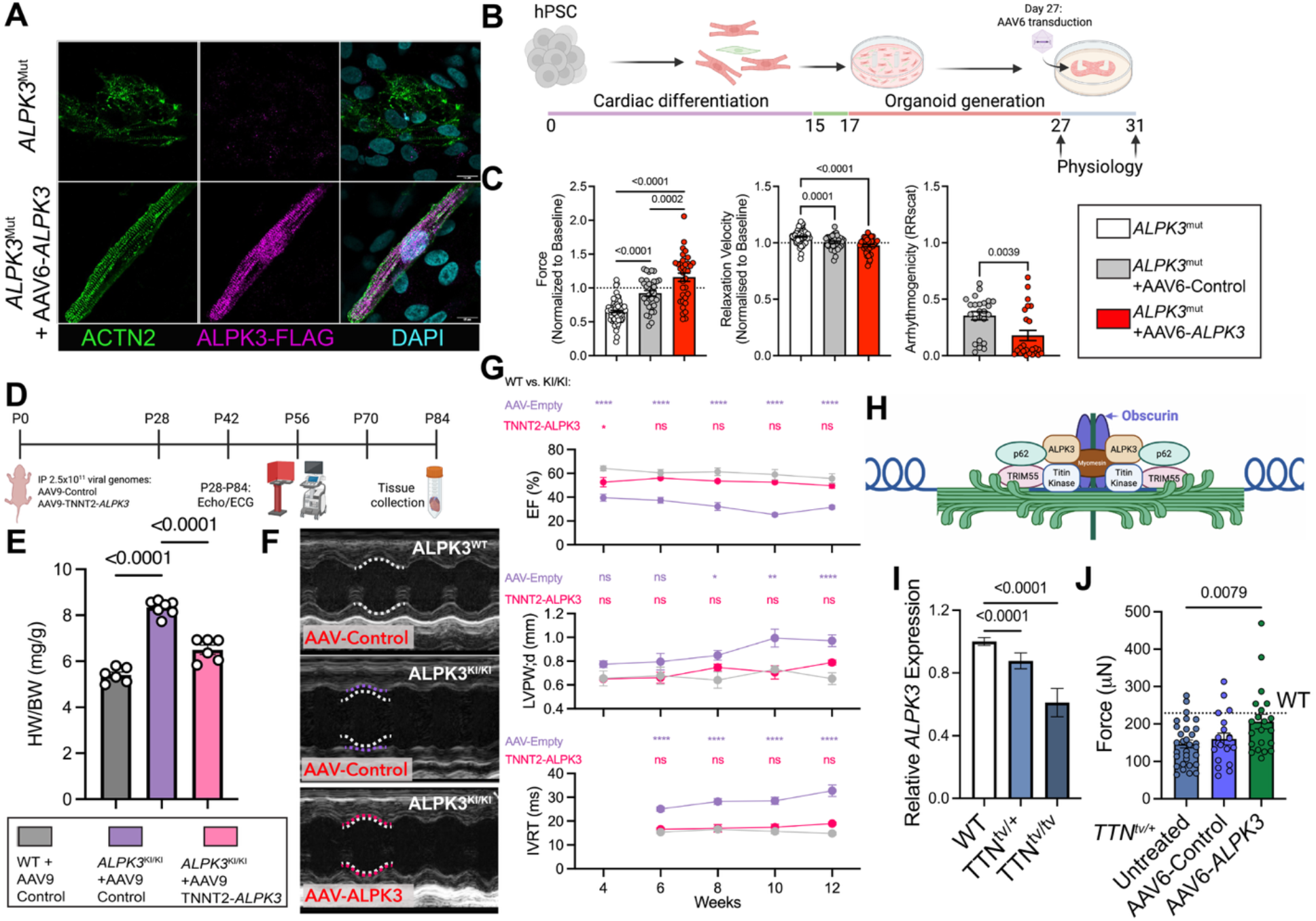
AAV-based *ALPK3* gene replacement therapy for genetic cardiomyopathy. (*A*) Delivery of AAV6-*ALPK3* to *ALPK3*^mut^ hPSC-derived cardiac monolayers demonstrated the correct localisation of ALPK3 to the M-Band and restore cardiomyocyte structure. (*B*) Schematic of HeartDyno^®^ experiments to test AAV-*ALPK3* restoration of function in *ALPK3*^mut^ cardiac organoids. (*C*) AAV6-*ALPK3* restored force deficits in *ALPK3*^mut^ cardiac organoids to the level of wild-type organoids (dotted line), without meaningful changes to contractile kinetics (n = 85 untreated, 32 AAV6-Control, and 34 AAV6-*ALPK3* individual organoids over three independent experiments; One-way ANOVA). AAV6-*ALPK3* also reduced arrhythmic events as measured by abnormalities in the peak-to-peak intervals (n=24 individual organoids over 3 independent experiments; unpaired t-test). (*D*) Experimental schematic for testing AAV9-*ALPK3* in CRISPR knock-in mice carrying the patient derived *ALPK3*^W153X^ variant. (*E*) Heart weight to body weight ratios for wild-type (WT) and *ALPK3*^W153X^ mice treated with AAV9-Control or AAV9-*ALPK3* demonstrated the restoration of heart mass to 63% of wild-type values (n=6 for WT, 7 for AAV9-Control, and 6 for AAV9-*ALPK3*; One-way ANOVA). (*F*) Representative parasternal long-axis M-Mode images for each group at 12 weeks-of-age. (*G*) Select echocardiography measurements (ejection fraction, diastolic posterior wall thickness, and isovolumic relaxation time) across the timescale of the experiment (n=6 for WT, 7 for AAV9-Control, and 6 for AAV9-*ALPK3*; Two-way ANOVA). (*H*) Graphical representation of the ALPK3/TTN protein quality control hub at the M-Band (*I*) Pseudobulk analysis of single nucleus RNA sequencing from an isogenic allelic series of hPSC-derived WT, *TTN*^tv/+^, and *TTN*^tv/tv^ cardiac cells demonstrated a dose-dependent reduction in *ALPK3* expression. (*I*) Heterozygous *TTN*^tv/+^ cardiac organoids exhibit reduced contractile function relative to isogenic WT organoids (dotted line). Transduction of *TTN*^tv/+^ cardiac organoids with AAV6-*ALPK3* restores this force deficit (n= 31 for untreated, 19 for AAV6-Control, and 21 for AAV-*ALPK3* from 4 independent experiments; ordinary One-way lognormal ANOVA).

Having confirmed cellular and structural rescue, we next assessed whether AAV6 mediated *ALPK3* expression could restore function in *ALPK3*^mut^ cardiac cells using the HeartDyno^®^organoid platform (Figure 1B).^6,9^ Consistent with our previous findings^6^, *ALPK3*^mut^ organoids exhibited reduced systolic force generation and increased propensity for asynchronous beating relative to isogenic controls (Figure 1C). We transduced organoids with 5×105 viral genomes per cell on day 27 and assessed function at baseline and 96 hours post-transduction. While AAV6-Control (*ALPK3* replaced with a non-coding sequence) could alter contractile force and kinetics, treatment with AAV-*ALPK3* restored active force production and relaxation kinetics to wild-type levels (Figure 1C). AAV6-*ALPK3* treatment also reduced asynchronous beating prevalence compared to AAV6-Control (Figure 1C).These findings demonstrate that AAV6-*ALPK3* efficiently rescues contractile function in a human model of *ALPK3* cardiomyopathy.

To validate these promising *in vitro* findings, we tested the efficacy of AAV9-*ALPK3* in an *in vivo* mouse model of *ALPK3* cardiomyopathy (Figure 1D). We used the *Alpk3*^W1538X^ mice which model a truncating variant found in patients.^2^ These homozygous mice are born with enlarged hearts, phenocopying the severity and early onset nature of this variant in humans.^2,6^ We delivered 2.5×10^11^ viral genomes of AAV9-*ALPK3* under the control of the cardiotropic *TNNT2* promoter via intraperitoneal injection at P0. AAV9-*ALPK3* demonstrated no adverse impacts on body weight or spleen mass in either group, despite a five-fold increase in *ALPK3* expression in the heart. By 12 weeks of age, treatment with AAV9-*ALPK3* significantly restored *Alpk3*^W1538X^ heart mass toward wild-type controls by 63% (Figure 1E). Biweekly echocardiography revealed near-complete restoration of all functional and structural parameters(Figure 1F-G), including ejection fraction (*Alpk3*^W1538X^ AAV9-Control 31±4% vs. *Alpk3*^W1538X^ *AAV9-ALPK3* 50±5%, p < 0.0001 at P84), stroke volume (*Alpk3*^W1538X^ AAV9-Control 27±4μL vs. *Alpk3*^W1538X^ *AAV9-ALPK3* 37±5μL, p = 0.0017 at P84; data not shown), wall thickness (diastolic posterior wall thickness: *Alpk3*^W1538X^ AAV9-Control 0.97±0.13mm vs. *Alpk3*^W1538X^ *AAV9-ALPK3* 0.79±0.05mm, p = 0.0341 at P84), and diastolic function (IVRT: *Alpk3*^W1538X^ AAV9-Control 32±7ms vs. *Alpk3*^W1538X^ *AAV9-ALPK3* 19±2ms, p < 0.0001 at P84). These results demonstrate that cardiotropic expression of full length *ALPK3* effectively restores heart function *in vivo* in *ALPK3* cardiomyopathy.

Mechanistically, we and others have discovered that ALPK3 orchestrates a protein quality control hub at the sarcomere to maintain turnover of contractile proteins.^6-8^ We therefore reasoned that AAV-*ALPK3* might have broader utility across other genetic cardiomyopathies that involve disrupted proteostasis^10^. We tested this hypothesis in an organoid model of *TTN* cardiomyopathy (*TTN*^tv/+^), which accounts for up to 25% of dilated cardiomyopathy cases. Importantly, the M-Band region of TTN shares an interaction hub with ALPK3 involving sarcomeric proteins as well as autophagy and proteasomal regulators at the M-Band of the sarcomere (Figure 1H).^6^ Single-nucleus RNA sequencing across an allelic series of *TTN*^tv/+^ hPSC-derived cardiomyocytes (*TTN* c.72663delA) revealed a gene dose-dependent impact of *TTN*^tv^ on *ALPK3* expression (Figure 1I). AAV6-mediated expression of full-length ALPK3 restored force production to wild-type levels in *TTN*^tv/+^ human cardiac organoids (Figure 1J), demonstrating the potential for an *ALPK3* gene therapy to address a broader spectrum of cardiomyopathies.

Herein we show that *ALPK3* gene therapy effectively restores contractile function in both human *in vitro* and *in vivo* mouse models of *ALPK3* cardiomyopathy. *ALPK3* cardiomyopathy represents up to 2% of hypertrophic cardiomyopathy cases^5^ – a target population comparable to established gene therapy target populations for Fabry disease or Friedreich ataxia. Moreover, our findings extend the potential of *ALPK3* gene therapy beyond this single indication due to the established role of ALPK3 in coordinating a protein quality control hub at the M-band of the sarcomere.^6^ Crucially, we demonstrate that *ALPK3* gene therapy restores function in cardiac organoids harboring *TTN*^tv/+^ variants, the most common genetic cause of dilated cardiomyopathy. This is particularly significant given that the extreme size of the *TTN* gene (>100,000 nucleotides) renders it unsuitable for conventional gene replacement therapy. These findings establish ALPK3 as a promising therapeutic approach for both *ALPK3* cardiomyopathy. It remains of interest to determine the range of TTN truncating variants, and potentially other cardiomyopathies with disrupted sarcomeric proteostasis, that may benefit from ALPK3 gene therapy.

## Acknowledgements

We thank the National Health and Medical Research Council of Australia (E.R.P., GNT2008376), the Heart Foundation of Australia (J.W.M., grant nos. 107192), the Medical Research Future Fund (J.W.M., E.R.P., D.A.E.; grant nos. MRF2024440), the Royal Children’s Hospital Foundation (E.R.P.) and the Murdoch Children’s Research Institute (MCRI) Early Career Researcher Award (J.W.M.) for grant and fellowship support. The MCRI is supported by the Victorian Government’s Operational Infrastructure Support Program. E.R.P., D.A.E., and R.J.M. are principal investigators of The Novo Nordisk Foundation Center for Stem Cell Medicine, reNEW, which is supported by Novo Nordisk Foundation grant no. NNF21CC0073729. The generation of the *Alpk3*^W1538X^ mice used in this study was supported by Phenomics Australia and the Australian Government through the National Collaborative Research Infrastructure Strategy program. We thank the Murdoch Children’s Research Institute iPSC Derivation and Gene Editing Facility for generation of the *TTN*^*tv*^ hiPSC lines.

